# Cryo-EM Reveals the Structural Basis of Microtubule Depolymerization by Kinesin-13s

**DOI:** 10.1101/206268

**Authors:** Matthieu P. M. H. Benoit, Ana B. Asenjo, Hernando Sosa

## Abstract

Kinesin-13s constitute a distinct group within the kinesin superfamily of motor proteins that promotes microtubule depolymerization and lacks motile activity. The molecular mechanism by which the kinesins depolymerize microtubules and are adapted to perform a seemingly very different activity from other kinesins is still unclear. To address this issue we obtained near atomic resolution cryo-electron microscopy (cryo-EM) structures of *Drosophila melanogaster* kinesin-13 KLP10A constructs bound to curved or straight tubulin in different nucleotide states. The structures show how nucleotide induced conformational changes near the catalytic site are coupled with kinesin-13-specific structural elements to induce tubulin curvature leading to microtubule depolymerization. The data highlight a modular structure that allows similar kinesin core motor-domains to be used for different functions, such as motility or microtubule depolymerization.

## Introduction

Kinesin-13s are a group within the kinesin superfamily of motor proteins that lack motile activity and instead work as microtubule depolymerases (Walczak et al., 2013). They are important regulators of microtubule dynamics in a variety of eukaryotic cell processes such as mitosis (Ganem and Compton, 2004; Manning et al.; Rogers et al., 2004), cytokinesis (Rankin and Wordeman, 2010), axonal branching (Homma et al., 2003) and ciliogenesis (Kobayashi et al., 2011) and are considered promising anti-cancer therapeutic targets (Ganguly and Cabral, 2015).

Like all kinesins, kinesin-13s posses a highly conserved ATPase motor-domain that it is essential for their function. The motor domain alone is sufficient to induce microtubule depolymerization (Asenjo et al., 2013; Moores et al., 2002; Tan et al., 2008) but an additional stretch of positively charged residues, N-terminal of the motor called the neck domain is necessary to achieve near full microtubule depolymerization activity (Hertzer et al., 2006; Maney et al., 2001).

The way that kinesin-13s interact with the microtubule is radically different from motile kinesins. Motile kinesins couple ATP hydrolysis to unidirectional stepping along the microtubule lattice. Kinesin-13s on the other hand couple ATP hydrolysis to the removal of tubulin subunits at the microtubule ends (Hunter et al., 2003) and when interacting with the microtubule lattice they undergo directional unbiased one-dimensional diffusion (Helenius et al., 2006). It is thought that kinesin-13s promote microtubule depolymerization by inducing a curved tubulin configuration incompatible with the microtubule tubular structure (Asenjo et al., 2013; Desai et al., 1999; Moores et al., 2002). Kinesin-13 constructs that include the motor domain form complexes with curved tubulin akin to the ones observed at microtubule-ends during depolymerization (Asenjo et al., 2013; Moores et al., 2003; Moores et al., 2002; Tan et al., 2006; Tan et al., 2008).

Although much is known regarding the mechanism of action of motile kinesins, how a highly conserved motor domain is adapted in kinesin-13s to interact with the microtubule in such a distinct manner, and induce microtubule depolymerization is still not clear. To address these questions we used cryo-electron microscopy to solve the cryo-EM structures of kinesin-13 constructs in complex with curved tubulin protofilaments or bound to the microtubule lattice.

## Results and Discussion

### KLP10A in Complex with Straight or Curved Tubulin

In the presence of ATP-like analogues KLP10A forms stable and very ordered complexes with curved tubulin. These complexes consist of curved tubulin protofilament with bound KLP10A and can be observed by electron microcopy in isolation, bound at the end of microtubules or wrapped around microtubules forming spirals (Asenjo et al., 2013). On the other hand in the absence of nucleotides (apo-state), KLP10A does not form complexes with curved protofilaments (Asenjo et al., 2013), but binds well to straight tubulin in the microtubule lattice. We solved the cryo-EM structures of minimal-MT-depolymerization-capable kinesin-13 constructs in complex with curved tubulin protofilaments or bound to the microtubule lattice (Fig. 1). The constructs based on the *Drosophila melanogaster* kinesin-13 KLP10A include either the motor-domain alone (M) or the motor plus the kinesin-13 specific neck-domain (NM: motor plus 81 residues N-terminus of the motor-domain). We solved the 3D structures of three complexes. The first one comprises the NM construct, bound to the microtubule (MT) lattice straight tubulin, and was prepared in the absence of nucleotides (NMMT_apo_, Fig. 1A). The second comprises the M construct, bound to curved tubulin (CT) protofilaments wrapped around microtubules, and was prepared in the presence of the non-hydrolysable ATP analogue AMP-PNP (CTMMT_AMP-PNP_, Fig. 1B). To determine the position of KLP10A neck domain in similar conditions we also performed 2D analysis of images of the NM construct in complex with curved tubulin protofilaments not wrapped around microtubules (Figs. 4 and S1D). The third complex comprises the NM construct, bound to straight tubulin on the microtubule lattice, and was prepared in the presence of AMP-PNP (NMMT_amp-pnp_, Fig. 1C). The overall resolution of the cryo-EM 3D density maps (3.5 to 4.1 Å) was sufficient to build atomic models in which the polypeptide chains were fully traced and the position of the ligands and many of the amino acid side chains determined (Figs. S2 and S3). Comparing the kinesin and tubulin structures in the three complexes revealed conformational changes associated with nucleotide binding and how they are coupled to the bound tubulin conformation.

**Fig. 1.**
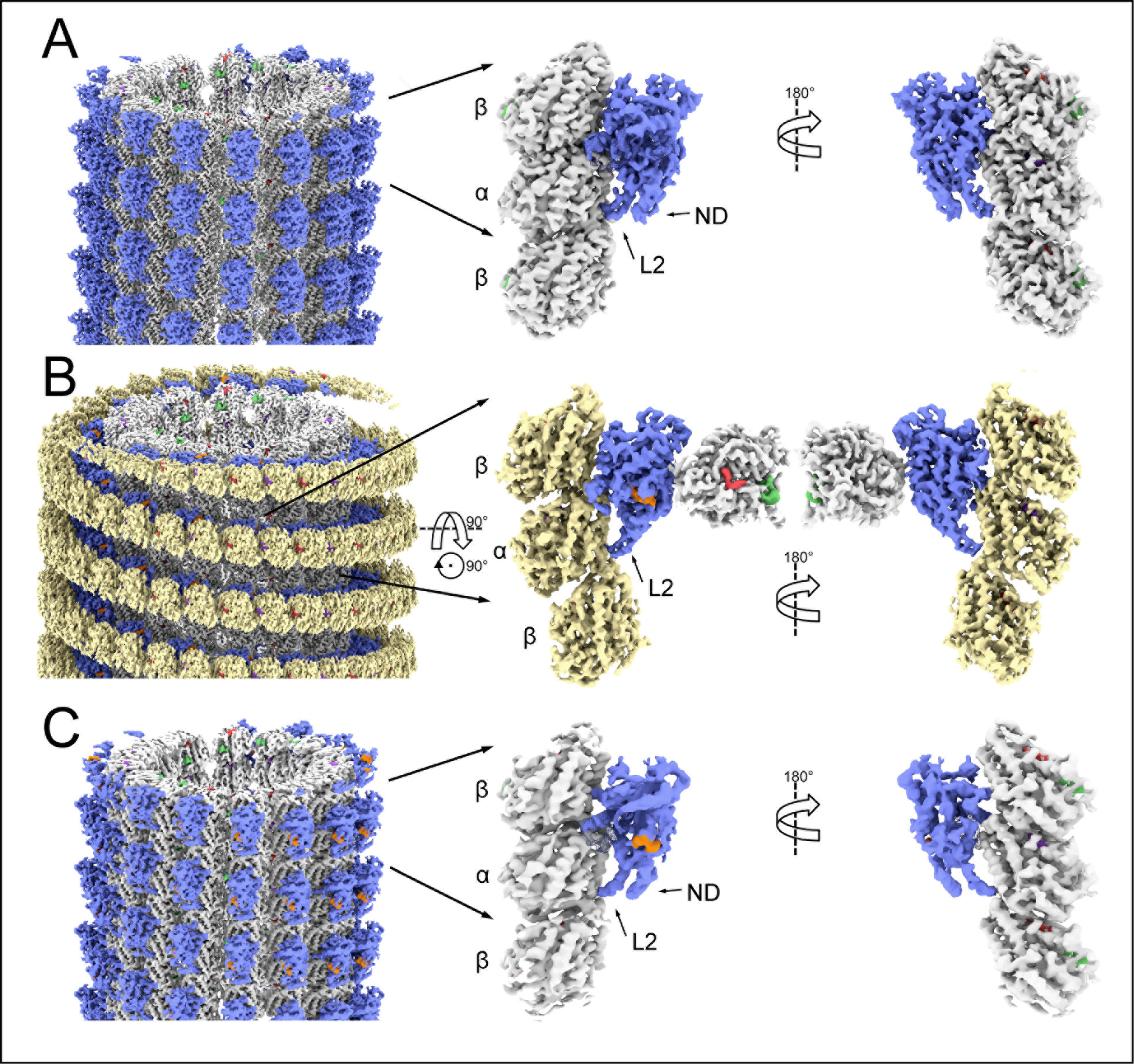
Cryo-EM 3D density maps. **(A)** NMMT_apo_ complex. **(B)** CTMMT_amp-pnp_ complex. **(C)** NMMT_amp-pnp_ complex. Left shows the whole helical 3D map, center and right shows two orientations of one asymmetric unit with an extra β-tubulin subunit. Coloring was done based on the fitted atomic models: microtubule lattice tubulin (straight tubulin) in white-grey, wrapping tubulin protofilament (curved tubulin) in ochre, KLP10A in blue, AMP-PNP in orange, GTP in purple. GDP in red, and paclitaxel in green. Note, that the interaction of the KLP10A motor-domain with straight tubulin in the NMMT complexes (A and C) and with curved tubulin in the CTMMT_AMP-PNP_ (B) complex are through the same putative kinesin-tubulin interface. An additional kinesin-13-specific tubulin binding site (Tan et al., 2008; Zhang et al., 2013) mediates the interaction of the motor domain with the microtubule in the CTMMT_amp-pnp_ complex.

### Nucleotide and Tubulin induced conformational changes in KLP10A

The structure of the KLP10A nucleotide binding site can be described in terms of the common structural elements found in other kinesins and the related NTPases, myosins and G-proteins (Kull and Endow, 2013; Vale, 1996). These elements include: the P-loop, a central β sheet with flanking α-helices and the switch loops (SW1 and SW2). The P-loop is involved in nucleotide binding and the switch loops sense the nucleotide species present in the active site by adopting distinct conformations (Vale, 1996). In the NMMT_apo_ complex, the kinesin ATP binding site is empty and the electron densities associated with switch loops regions are positioned relatively far away from the nucleotide site (Figs. 2A and 2D). In analogy with other NTPases this corresponds to the ‘open’ configuration of the catalytic site. In contrast, in the CTMMT_AMP-PNP_ complex the KLP10A nucleotide binding pocket adopts the ‘closed’ configuration. A clear electron density associated with the bound AMP-PNP is observed (Figs. 1B, 2B and 2E), the switch loops are closer to the nucleotide γ-phosphate and α-helix 4 (H4) which is part of SW2 adopts a different orientation relative to the rest of the motor-domain (Fig. 2A-F, Movie S1).

**Fig. 2.**
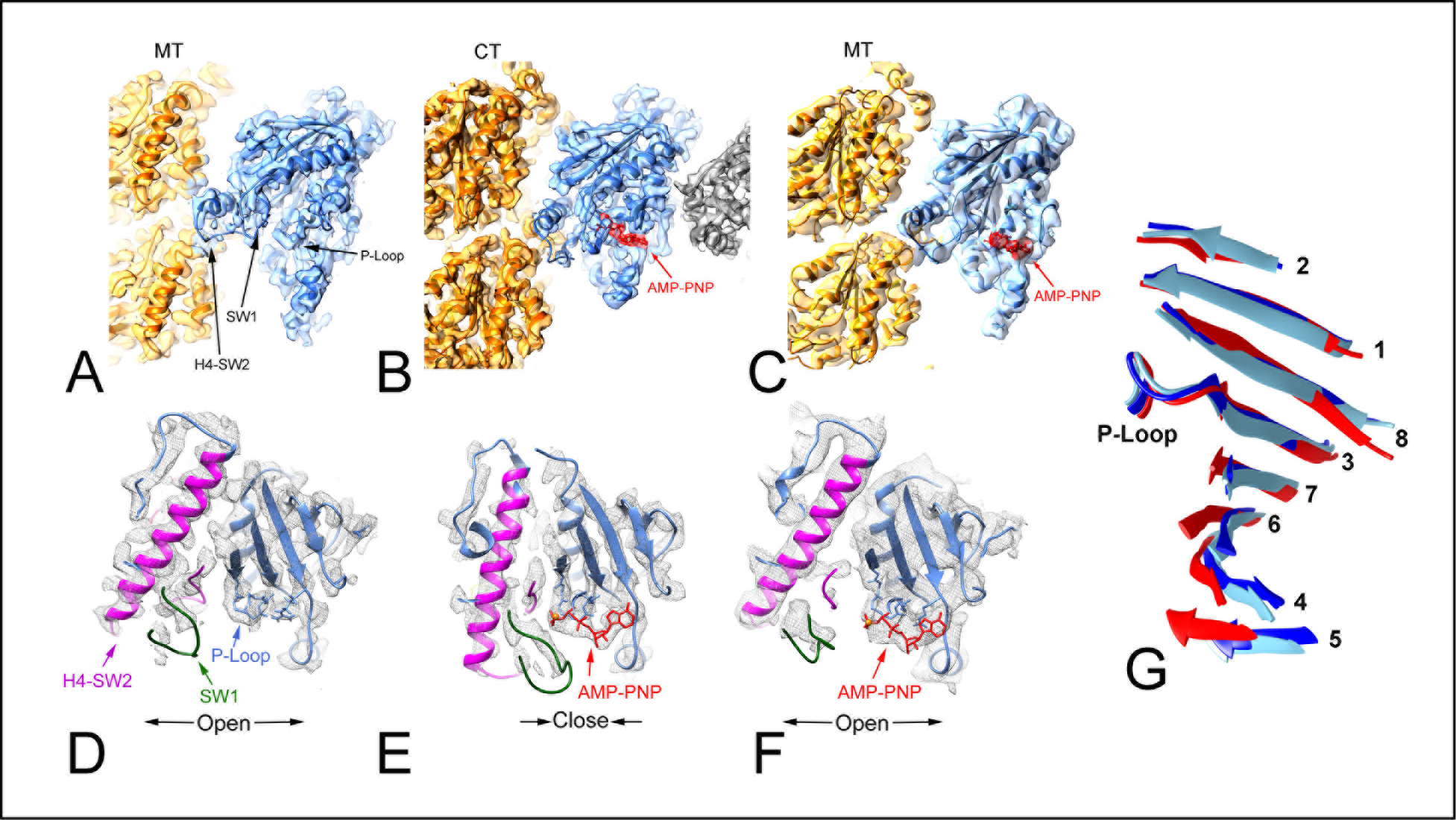
KLP10A nucleotide binding pocket. **(A-C)** Side views (tubulin plus end up) showing the KLP10A nucleotide binding pocket of the complex models. **(A)** NMMT_apo_. **(B)** CTMMT_AMP-PNP_. **(C)** NMMT_AMP-PNP_. **(D-F)** Top view detail of the KLP10A nucleotide binding pockets. **(D)** NMMT_apo_. **(E)** CTMMT_AMP-PNP_. **(F)** NMMT_AMP-PNP_. **(G)** P-loop aligned central β-sheets of the 3 complexes, NMMT_apo_ in blue, CTMMT_AMP-PNP_ in red and NMMT_AMP-PNP_ in light blue. Numbers indicate the strand order in the KLP10A sequence.

The closed configuration of the nucleotide binding pocket found in the CTMMT_AMP-PNP_ complex is similar to the one found in the crystal structures of motile kinesins thought to represent the hydrolysis competent ATP bound form (Parke et al., 2010). It is also similar to the crystal structure reported recently of another kinesin-13 (hMCAK) in complex with curved tubulin (Wang et al., 2017). These similarities indicate that the nucleotide pocket of kinesin-13s and motile kinesins undergo similar conformational changes in response to the nucleotide specie in the active site. However, our results also indicate that in kinesin-13 these conformational changes are coupled to the nucleotide specie and to the particular tubulin configuration, straight or curved to which the motor domain is bound. The nucleotide pocket is closed with AMP-PNP only when bound to curved tubulin and it is open when bound to straight tubulin, independently of the nucleotide species in the active site (Fig. 2A-F).

Near the nucleotide binding pocket we also observed structural differences between KLP10A and other kinesins that are likely related to the distinct functionality of kinesin-13s. Loop-5, near the P-loop, is in a different configuration than the one observed in other kinesins, where it is either not resolved or it is in a different configuration than the one observed here (Fig. S4). Residues in this area are not highly conserved between kinesin families and it is thought that distinct L5s modulate the ATPase activity of different kinesins (Muretta et al., 2013). Accordingly, the distinct configuration observed here may be partially responsible for some of the unique ATPase properties of kinesin-13s (Friel and Howard, 2011; Wang et al., 2012).

Further away from the nucleotide pocket the switch region movements detected between the NMMT and the CTMMT_AMP-PNP_ complexes, are propagated through other regions of the motor domain. The amount of twist of the central β-sheet is different (Fig. 2G). Also the reorientation of H4, which is part of the kinesin-tubulin interface in the complexes, causes a reorientation of the rest of the motor domain relative to the bound tubulin. The ultimate consequence of all these structural differences is a reshaping of the tubulin binding interface so that it alternatively complements the shape of straight or curved tubulin (Fig. 3). From the KLP10A structure in the NMMT_apo_ or NMMT_AMP-PNP_ complexes to the one in the CTMMT_AMP-PNP_ complex, loop-2 (L2) moves relative to the other areas of the kinesin-tubulin interface (Fig. 3B). L2 interacts with the tubulin intradimer interface and the interaction is maintained in the different complexes by a concomitant tubulin conformational change from the straight to the curved configuration (Fig. 3C, Movies S4 and S5). The interacting residues comprise the kinesin-13 class conserved motif KVD at the tip of loop-2 and complementary residues located in α- and β- tubulin at the interdimer interface (Fig. 3A). This interaction is critical for kinesin-13 function as mutating the residues KVD to alanines abolishes kinesin-13 microtubule depolymerase activity (Ogawa et al., 2004; Shipley, 2004; Tan et al., 2008). The movement of L2 relative to the rest of the kinesin tubulin interface is analogous to the power stroke occurring in motile proteins, where a change in the nucleotide species in the active site causes a rearrangement of the switch loop regions. This conformational change ultimately provokes a change in the position or orientation of an ‘amplifier’ element such as the lever arm in the case of myosins, or the neck/neck-linker in the case of motile kinesins (Kull and Endow, 2013; Shang et al., 2014).

**Fig. 3.**
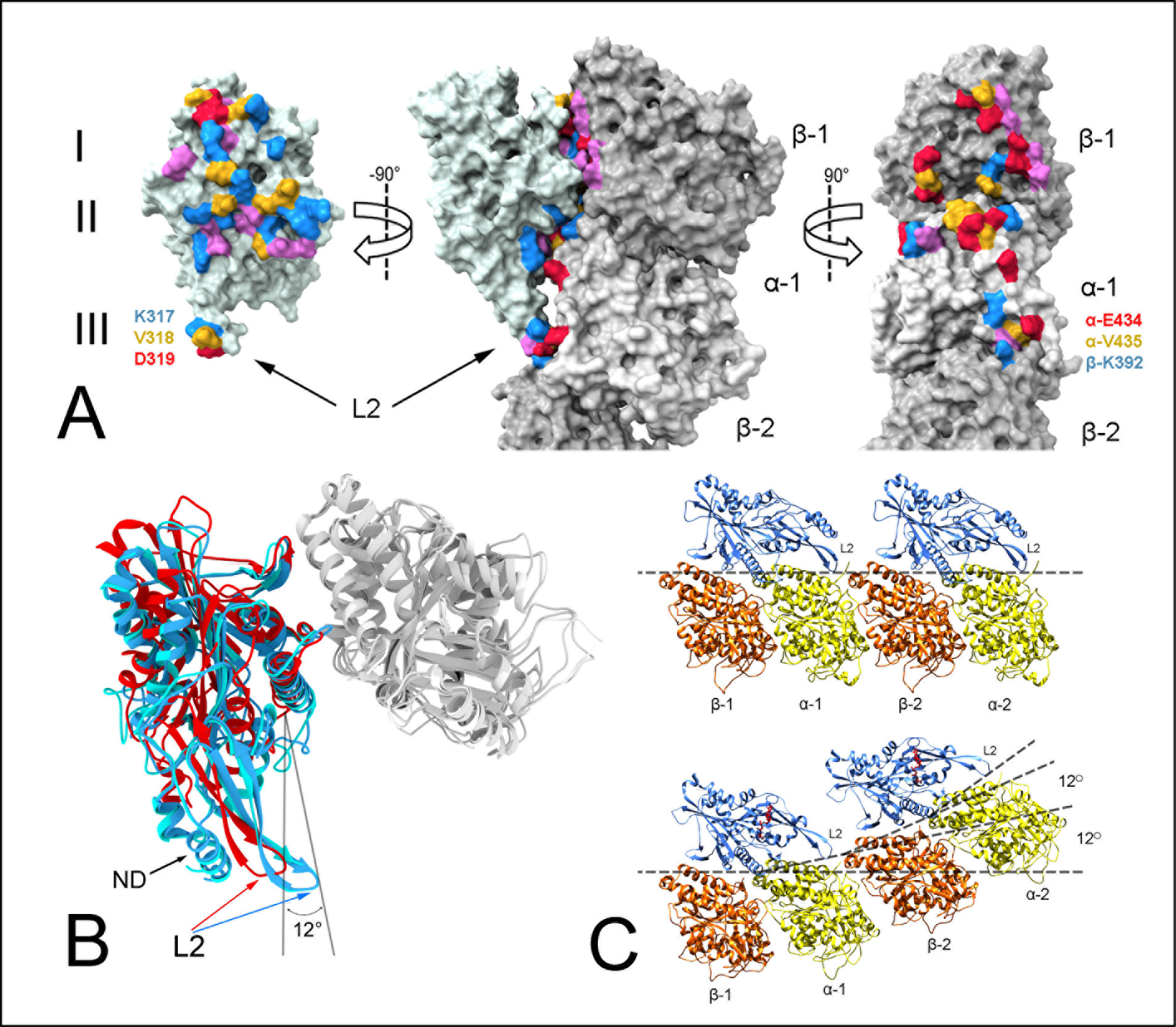
KLP10A-tubulin binding interface. **(A)** Solvent accessible surface of the CTMMT_AMP-PNP_ complex highlighting residues at the tubulin KLP10A interface within contact distance (NMMT_apo_ and NMMT_AMP-PNP_ complex interfaces are shown in Fig. S5). The interface can be divided in three areas. Area I towards β-tubulin plus side, area II near the tubulin intra-dimer interface and area III near the inter-dimer interface. Residues are color coded according to type. Negative red, positive blue, polar magenta and non-polar yellow. **(B)** Superimposed KLP10A model structures aligned on the bound β-tubulin subunit (and interface area I). Red: CTMMT_AMP-PNP_ complex; Blue: NMMT_apo_; Light blue: NMMT_AMP-PNP_. **(C)** Side views of a protofilament of the NMMT_apo_ (top) and CTMMT_AMP-PNP_ (bottom).

### Conformation and Position of the KLP10A-Neck-Domain

The KLP10A M constructs without the neck-domain bind to every tubulin heterodimer along the protofilament as shown in the CTMMT_AMP-PNP_ complex. However, constructs that include the neck-domain can be found binding to every tubulin heterodimer (Fig. 4B, bottom row) or consistently skipping one tubulin heterodimer (Fig. 4B, top row). This second binding pattern results in a stoichiometry of one KLP10A molecule per two tubulin heterodimers. A similar tubulin decoration pattern was observed previously with another kinesin-13 construct that also included the neck-domain (Mulder et al., 2009) and suggests that the neck domain is positioned in such a way to prevent binding of and adjacent motor domain. Class average images of complexes with this pattern of one KLP10A NM construct per two tubulin heterodimers reveals an elongated density going from the back of the motor domain to the intradimer interface of the next tubulin heterodimer (Fig. 4B top row). This elongated density can be well fitted to an α-helical structure consistent with the predicted secondary structure of the neck-domain sequence (Fig. 4C). Part of the neck domain is resolved forming a loop and an α-helix at the back of the motor domain in the NMMT complexes 3D structures. (Fig. 1A, 1C, 3B). These results indicate that the kinesin-13 neck domain consists mainly of an α-helical structure that provides additional kinesins-13 interactions with the next tubulin heterodimer along the protofilament. This configuration of the neck domain is consistent with its proposed role in assisting kinesin-13 microtubule binding (Chatterjee et al., 2016; Cooper et al., 2010; Mulder et al., 2009). It may also help to stabilize tubulin inter-dimer curvature. However, the neck domain is unlikely to be the major driver of tubulin curvature as the M construct can readily form stable complexes with curved tubulin.

**Fig. 4.**
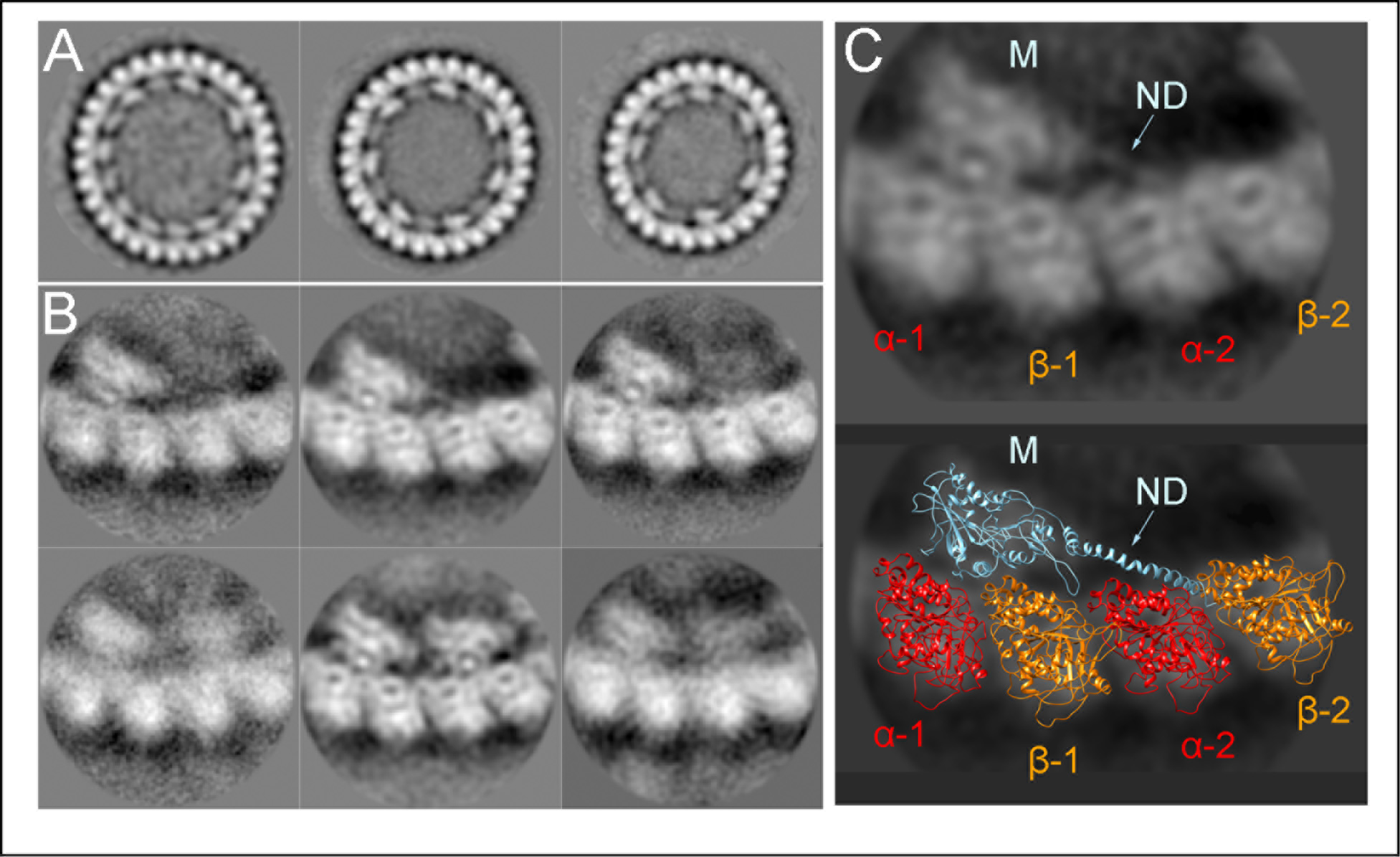
Curved tubulin KLP10A-Neck motor AMP-PNP complex (CTNM_AMP-PNP_) (A) Class averages of curved tubulin protofilaments obtained by aligning/classifying the whole ring-like structures observed in cryo-EM images (Fig. S1D). (B) Class averages of curved tubulin protofilaments obtained by aligning/classifying regions containing two tubulin heterodimers from the classes above. This classification produced classes in which only one KLP10A motor domain was bound to two tubulin heterodimers (top) and others in which each of the two tubulin heterodimers had one KLP10A motor domain bound (bottom). (C) In the class average images with only one KLP10A motor domain per two tubulin heterodimers, an elongated density can be observed going from the motor domain density (M) to the density corresponding next tubulin heterodimer (α-2, β2). This elongated density is modeled as an α-helix formed by the KLP10A neck-domain (ND).

### Tubulin conformational changes

The structural differences between straight and curved tubulin can be described to a first approximation as rigid body rotation of 12° between tubulin subunits, or 24° per tubulin heterodimer (Fig. 3B and 3C). These are the values obtained from the structure of the wrapped around protofilament of the CTMMT_AMP-PNP_ complex (helical path with 23.82° rotation and 11.1Å rise per tubulin dimer). In KLP10A-tubulin complexes not wrapped around microtubules a range of curvatures were observed up to 27° per tubulin heterodimer (Fig. 4A). In addition to this rotation between monomers, there are other small structural changes within the tubulin subunits. The root mean squared deviation (RMSD) between equivalent α-carbon positions of straight and curved tubulin were 1.9 Å and 2.0 Å for the α- and β-tubulin subunits respectively. A conformational change to highlight is the disruption of the binding site of the anti-cancer drug paclitaxel (Taxol^®^) on the β-subunit of KLP10A bound curved tubulin. Despite all complexes being made in the presence of paclitaxel (see Methods) the corresponding density is absent in the curved tubulin and the binding site is partially blocked by β-tubulin residues (Fig. S6). In contrast, a paclitaxel associated density is present in the microtubules of all three complexes investigated (Fig. 1). This includes the microtubule in the CTMMT_AMP-PNP_ complex that also contains the sans-paclitaxel curved tubulin.

### A pre-power stroke configuration

The structure of the NMMT_AMP-PNP_ complex provided further insights into the relationship between the nucleotide species in the kinesin-13 active site and tubulin conformation. Surprisingly, in this complex the KLP10A switch loops remain in the open configuration despite the presence of AMP-PNP in the active site (Figs. 2C and 2F). The structure is closer to the apo state than to the one in the presence of AMP-PNP, but bound to curved tubulin (Figs. 2, 3B, Movies S2, S3, S5 and S6). This result indicates that the KLP10A conformational changes associated with ATP binding (as mimicked by AMP-PNP) are prevented when bound to straight tubulin in the microtubule lattice. In analogy with motile motor proteins this structure would represent a “pre-power stroke” configuration; where the nucleotide species promoting the power stroke is in the active site, but movement of the amplifier element is mechanically prevented. In addition, given that ATP hydrolysis is thought to be catalyzed by the closed configuration of the nucleotide pocket (Kull and Endow, 2013; Parke et al., 2010), the structure implies that KLP10A is not catalytic when bound to the microtubule-lattice trough the putative kinesin-tubulin binding site. Only the KLP10A structure bound to curved tubulin in the presence of AMP-PNP (CTMMT_AMP-PNP_ complex) has the nucleotide pocket in the closed configuration while it remains open when bound to straight tubulin (NMMT_apo_ and NMMT_AMP-PNP_ complexes), regardless of the presence of nucleotide. It is reported that the ATPase activity of kinesin-13s are stimulated to a higher level by interaction with the microtubules ends in comparison with interaction with the microtubule lattice (Hunter et al., 2003). Our structural results strongly support a model in which this differential ATPase stimulation is the result of an acceleration of the hydrolysis step in the complex formed with curved tubulin at the microtubule ends (Fig. 5).

**Fig. 5.**
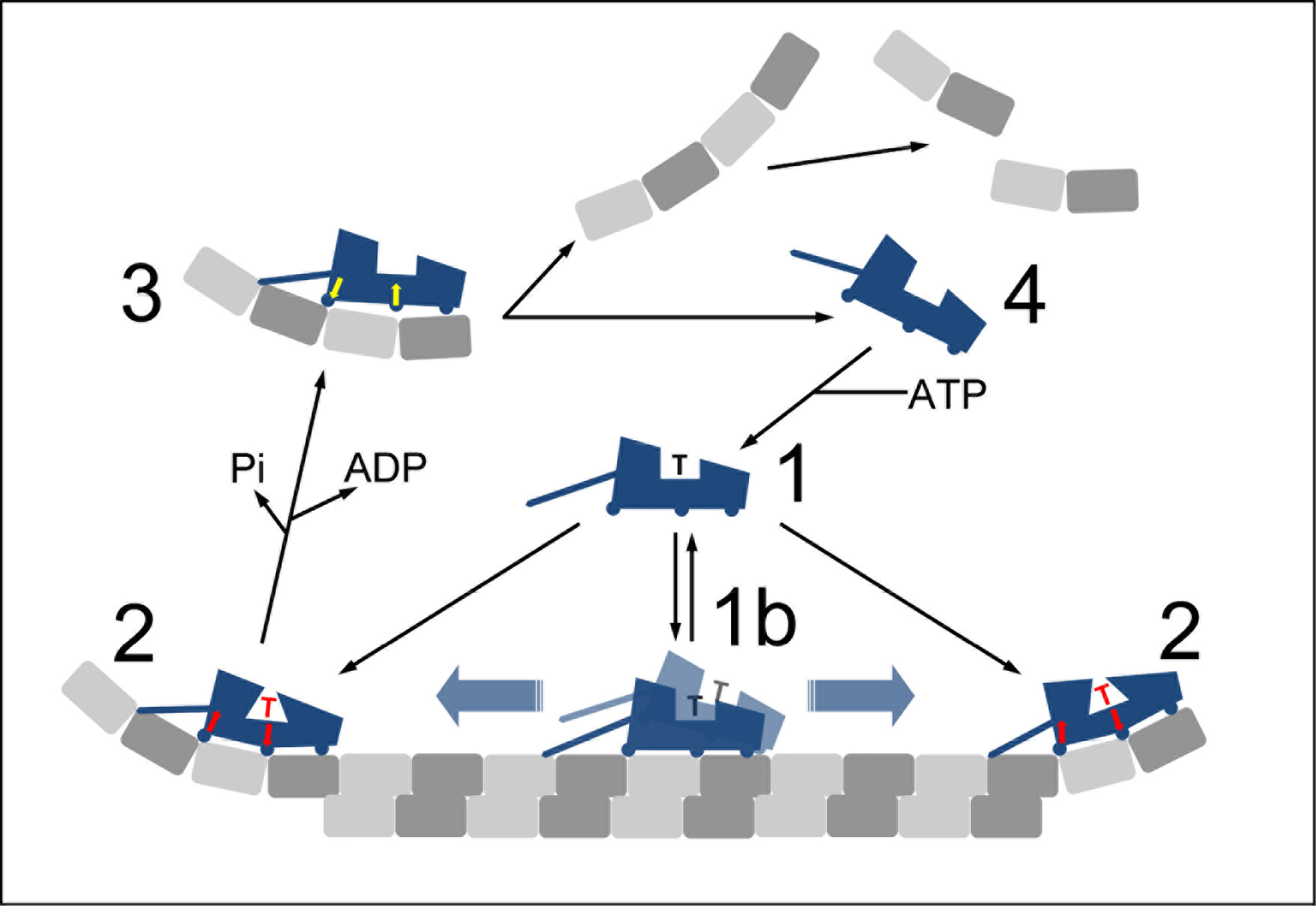
Kinesin-13 microtubule depolymerization model. **State 1**: When not interacting with tubulin, kinesin-13s are predominantly in an ATP bound form (Friel and Howard, 2011; Hunter et al., 2003; Wang et al., 2012). State 1b: Binding to the microtubule lattice results in a relatively weak interaction (Wang et al., 2012) where the kinesin motor domain can detach from the microtubule or remain attached in a non-stereo specific-highly-mobile mode that can diffuse along the microtubule lattice (Chatterjee et al., 2016) (one-dimensional diffusion). In this state our structural results predict that kinesin-13 stays predominantly in an ATP bound form as the nucleotide pocket is in the open configuration (non-catalytic) when bound to the microtubule lattice (Fig. 2). Alternatively, interaction with the microtubule lattice could allow closure of the binding pocket (e.g. during microtubule unbinding) leading to ATP-hydrolysis and to the diffusive ADP-Pi bound form proposed to exists in other models (Friel and Howard, 2011; Helenius et al., 2006). **State 2:** An ATP containing kinesin-13 binds and stabilizes curved protofilaments already present at the microtubule ends or bends tubulin protofilaments lacking a full set of lateral contacts. In either case, binding to curved tubulin enables closure of the nucleotide pocket promoting ATP hydrolysis (indicated by the arrows, the triangular shape of the binding pocket, and the red letter T). **State 3:** After hydrolysis and product release, the motor domain alters its shape to an apo configuration that it is no longer complementary to the curved tubulin leading to detachment from curved tubulin. We propose that this occur after product release as curved-tubulin-KLp10A complexes can form in nucleotide conditions mimicking the ADP-Pi and ADP states but not the apo state (Asenjo et al., 2013). **State 4:** After kinesin detachment the curved protofilaments loose potentially stabilizing inter-dimer interactions with kinesin-13 loop-2 and dissociate into smaller oligomers. Binding of ATP to the dissociated motor domain completes the cycle (State-1).

In summary, our structural studies provide an atomic level detail of the mechanism of microtubule depolymerization by kinesin-13 and how it is coupled to ATP hydrolysis. They show how kinesin-13 binding alters tubulin structure and conversely how tubulin binding alters the structure of the kinesin-13 ATP binding site.

## Materials and Methods

Protein expression. Kinesin proteins were bacterially expressed in BL21 cells (BL21(DE3)pLysS, Thermofisher). The motor construct (M) includes *Drosophila melanogaster* KLP10A residues 279-615 and was produced as previously reported (Tan et al., 2006). The neck motor construct (NM) includes KLP10A residues 198-615 and was produced as follow: KLP10A residues 198-615 were cloned in a pRSET B plasmid with an N-terminal histidine tag. A TEV protease site (ENLYFQG) was introduced between the His tag and the start of the KLP10A sequence. Plasmid transformed cells were grown (6 to 8 l) in LB media containing 200 μg/ml ampicillin and 34 μg/ml chloramphenicol. The cultures were inoculated at an OD_600nm_ of 0. 2 to 0.3 from an overnight preculture in the same media. One liter culture in 2l flasks in the same media were incubated at 240 rpm, 37°C until the OD_600nm_ reached 1.2. Protein expression was induced by adding 700 μM IPTG and incubating for 7 hours at 30°C and 200 rpm. Cells were spun for 15 min at 3000 × g and the pellet stored at - 80°C. All the purification steps below were performed at 4°C. Bacteria cells were lysed at 4°C with a microfluidizer in buffer A (1 mM MgCl_2_, 1 mM EGTA, 250 mM KCl, 1 mM Mg-ATP, 5 mM β-mercaptoethanol, 80 mM K-PIPES, pH 7.2, adjusted with KOH) supplemented with 10 mM imidazole and one pellet of anti-protease cocktail (Sigmafast Protease Inhibitor Cocktail Tablets, EDTA-Free, Sigma-Aldrich) per 20 g of cell paste. The lysate was spun for 40 min at 150,000 × g and 4°C. The supernatant was passed through a NiNTA column with 5 ml of resin pre-equilibrated with buffer A supplemented with 10 mM imidazole. The column was washed sequentially with 100 ml of buffer A supplemented with 50 mM imidazole, 100 ml of buffer A with 1 M KCl and 8 ml buffer A. Elution was performed by adding 1 ml of buffer A supplemented with 300 mM imidazole.

Fractions containing the proteins were pooled and subjected to dialysis overnight in buffer A with histidine-tagged TEV protease (Harder et al., 2008) to cleave the tag (TEV/kinesin molar ratio of 1/20). TEV protease was expressed in transformed BL21(DE3)-RIL cells with plasmid pRK793 (Kapust et al., 2001) (Addgene plasmid # 8827, courtesy of David Waugh lab).

After dialysis, the protein solution was passed 1 ml at a time on a 5 ml NiNTA column preequilibrated with buffer A supplemented with 20 mM imidazole and the eluted fraction were collected. The untagged neck-motors in the column were released by washing the column – 1 ml at a time – with a total volume of 20 ml of the equilibration buffer supplemented with 20 mM imidazole. The collected fraction containing the (untagged) neck-motor construct were pooled, concentrated to 600 μl and loaded on a Superdex 200 Increase 10/300 GL column preequilibrated with buffer B (1 mM MgCl_2_, 1 mM EGTA, 250 mM KCl, 150 μM Mg-ATP, 5 mM μ-mercaptoethanol, 80 mM K-PIPES, pH 7.2). The purified kinesin solution was concentrated, supplemented with 20 % (v/v) sucrose, aliquoted and flash frozen at −80°C with a final protein concentration of 140 to 150 μM. KLP10A concentration was estimated by UV absorbance at λ=280nm using a Nanodrop (ThermoFisher, MA) and an extinction coefficient of 23300 L.mol^−1^. cm^−1^ (value estimated from the protein sequence and assuming one ATP molecule bound).

**Microtubules**. Microtubules were prepared from porcine brain tubulin (Cytoskeleton, Inc. CO). Tubulin lyophilized pellets were resuspended in BRB80 (80 mM K-PIPES, 1 mM MgCl_2_, 1 mM EGTA, pH 6.8) to 45 μM and spun at 313,000 × g before polymerization to eliminate aggregates. Microtubule polymerization was done in conditions to enrich the number of microtubules with 15 protofilaments (Wilson-Kubalek et al., 2016) as follow. The clarified resuspended tubulin solution was supplemented with GTP, MgCl_2_, DMSO to final concentrations of ~35 μM tubulin, 80 mM PIPES, pH 6.8, 1 mM EGTA, 4 mM MgCl_2_, 2 mM GTP, 12% (v/v) DMSO and incubated 40 minutes at 35°C. An aliquot of stock Paclitaxel (Taxol®) solution (2 mM in DMSO) was added to a final paclitaxel concentration of 250 μM and incubated for another 40 minutes at 35°C. The microtubules were then spun at 15,500 × g, 25°C and the pellet resuspended in BRB80 with 20 μM paclitaxel. A slightly different microtubule polymerization solution (80 mM PIPES, pH 6.8, 1 mM EGTA, 4 mM MgCl_2_, 2 mM GTP, 20 μM paclitaxel) was used to image NM-curved tubulin-protofilament complexes not wrapped around microtubules (CTNM_AMP-PNP_ complex, Fig. 4). Tubulin concentration was estimated by UV spectroscopy using an extinction coefficient of 115000 L·mol^−1^·cm^−1^ on a solution of microtubule with Guanidine Hydrochloride at a final concentration of 5.25 M.

### Cryo-EM sample preparation

NMMT complexes. NM aliquots were thawed on ice and buffer exchanged at 4°C to BRB40 (40 mM K-PIPES, pH 6.8, 1 mM MgCl_2_, 1 mM EGTA) supplemented with 50 μM ATP, using a spin-column (Micro Bio-Spin 6 Column, Bio-Rad) and kept on ice.

A solution containing the NM and the appropriate nucleotide conditions to be tested (NM mix) was prepared: 88 μM NM construct in BRB40 supplemented with 20 μM paclitaxel and either 20.10^−3^ unit of potato apyrase (Sigma-Aldrich, MO) for the apo state or 4 mM AMP-PNP (Sigma-Aldrich, MO) and 4 mM MgCl_2_ for the AMP-PNP state (all final concentrations). Four micro-liters of a microtubule solution (8 μM tubulin in BRB80 plus 20 μM paclitaxel) were layered onto untreated EM grids (Quantifoil carbon/copper grids 300 mesh R2/2), incubated 1 minute at room temperature and then the excess liquid removed from the grid using a Whatman #1 paper. Four microliters of the NM mix were then applied onto the EM grid and incubated for 1 min at room temperature and then the excess liquid blotted as before. A second application of 4 μL of a freshly prepared NM mix to the grid was performed and then the grid was mounted into a Vitrobot apparatus (FEI- ThermoFisher MA), incubated 1 minute at room temperature and plunge frozen into liquid ethane. Vitrobot settings: 100% humidity, 3 seconds blotting with Whatman #1 paper and - 2 mm offset. Grids were stored in liquid nitrogen until imaging in a cryo-electron microscope.

**CTMMT_AMP-PNP_ complex**. Microtubules and KLP10A constructs where mixed into an incubation solution: 3 μM KLP10A M construct, 1.5 μM polymerized tubulin in BRB80, with 2 mM AMP-PNP, 2 mM MgCl_2_ and 20 μM paclitaxel (all final concentrations). The mix was incubated at room temperature for 20 min and spun for 10 min at 28°C at 30,000 × g. The pellet was resuspended in one fifth of the initial volume. Four microliters of the resuspended solution were layered into an untreated electron microscope grid (Quantifoil carbon/copper grids 400 mesh R2/4) and incubated 1 minute at room temperature. The grid was then placed into a Vitrobot apparatus for plunge freezing into liquid ethane. Vitrobot settings: 100% humidity, 2 seconds blotting with Whatman #1 paper and - 2 mm offset.

**CTNM_AMP-PNP_ complex.** NM aliquots were thawed on ice and their storage buffer was exchanged at 4°C to BRB80 supplemented with 50 μM ATP using a spin-column (Micro BioSpin 6 Column, Bio-Rad) and stored on ice. Microtubules and the KLP10A NM construct were combined into an incubation mix: 15 μM KLP10A NM construct and 3 μM polymerized tubulin in BRB80 buffer supplemented with 5 mM AMPPNP, 5 mM MgCl_2_ and 20 μM paclitaxel. The mix was incubated 5 minutes at room temperature and then layered into untreated electron microscope grids and plunge frozen as explained above for the NMMT complexes.

**Cryo-EM data acquisition.** Data were collected at 300 kV on a Titan Krios microscope equipped with a K2 summit detector. Acquisition was controlled using Leginon (Suloway et al., 2005) with the image-shift protocol. Data collection was performed manually. For each hole selected, a picture at low magnification (2692X) was used to target 1 to 3 filaments on which image-movies where taken. Image-movies consisted of 50 to 60 frames of 200 ms/frame and 1.24-1.28 e^−^ /Å exposure per frame. For the microtubule decorated complexes image-movies were collected within a nominal defocus range of −0.5 to −1.5 μm. This resulted in a distribution of image defocus of −1.6 ± 0.8 μm (median ± SD) as estimated with Gctf (Zhang, 2016). Individual NM-curved tubulin complexes were imaged at a higher defocus (−3.0 ± 0.5 μm).

**Image analysis and 3D reconstruction.** Movie frames were aligned with motioncor2 (Zheng et al., 2017) (v01-30-2017) using default parameters, generating both dose weighted and non dose weighted sums. CTF parameters per micrographs were estimated with Gctf (Zhang, 2016) (v0.50) on aligned and non dose weighted movies average.

The number of cryo-EM images and asymmetric units (au) processed for the 3D reconstructions where as follow: NMMT_apo_ complex, 155958 au from 348 images; CTMMT_AMP-PNP_ complex, 242762 au from 560 images; NMMT_AMP-PNP_ complex, 88909 from 584 images (1 au corresponding to one microtubule tubulin dimer and associated proteins).

A helical-single-particle approach was used to obtain 3D electron density maps of microtubules decorated with KLP10A neck-motor construct (NMMT complexes) and with KLP10A motor construct-curved tubulin protofilament spirals (CTMMT complex). The approach consisted of the following steps:

Step 1. Microtubules were manually picked on the aligned micrograph averages using a custom script written in the R programming language.

Step 2. Microtubule symmetry identification and selection. Atomic models of microtubules ranging from 11 to 16 protofilaments of type R of L (right or left handed protofilament twist) were generated, and converted to densities with EMAN1 (Ludtke et al., 1999) pdb2mrc with a voxel step size of 4 Å/voxel. The densities were low passed filtered to 15 Å and the Spider (Frank et al., 1996) operation PJ 3Q was used to generate 200 reference projections per microtubule type. Experimental microtubule image segments were extracted and resampled to 4 Å/pixel. Segment corresponding to each experimental microtubule image were subjected to multi-reference alignment and classification with the set of model microtubule image projections (Spider (Frank et al., 1996) AP SHC operation) to identify the best matching pairs. A microtubule type was assigned to each experimental filament image as the microtubule type with more matches to the experimental filament image. Only filaments with a majority of the corresponding segments identified as 15R (15 protofilaments with right handed supertwist) were selected for further processing. This procedure was repeated using as reference a 15R microtubule reconstruction generated from the experimental data as explained in step 3. At the end only microtubule segments with of at least 10 successive particles identified as 15R were retained for further processing.

Step 3. Initial model. To get an initial model a first round of 3D reconstruction was performed following the procedure described in step 5 but using a single reference map instead of two independent ones. The reference maps were generated by converting to electron densities the PDB model 3J2U for the CTMMT complex or a modified version with the kinesin motor directly docked on the microtubule for the NMMT complexes. The PDBs atomic coordinates were converted to electron densities using EMAN1 (Ludtke et al., 1999) pdb2mrc and low-pass filtered to 20 Å. After this first 3D reconstruction the helical symmetry parameters of the resulting maps were determined using the Relion 2.0 helix toolbox (He and Scheres, 2017). The estimated parameters, rotation (ϕ) and rise (r) per subunit were ϕ=168.083°, r=5.50 Å for the NMMT complexes and ϕ=168.089, r=5.57 Å for the CTMMT complex.

Step 4. Microtubule axis refinement. To limit the allowed translations of the particles in the 3D refinement reconstruction procedure (step 5) we used an iterative procedure to refine the coordinates of the microtubule axis. First the manually picked microtubules coordinates were fitted by a spline curve and interpolated to about 15 times the helical rise (82 Å). These coordinates were used to extract 300 × 300 pixels 2X binned, boxed particle images (low-pass filtered to 15 Å) which were subjected to 4 centering cycles. In each cycle the particles were aligned using the Spider (operation AP SHC) against projections of the initial model created in step 3. From the translation values obtained from the projection matching procedure a new set of axis coordinates was determined. This set was fitted to a spline curve with a maximum degree of freedom of 2, and re-interpolated every 82 nm along the axis to generate a new set of coordinates for the next cycle (4 cycles).

Step 5. Position-orientation refinement and 3D reconstruction. Refinement was done completely independently in two halves of the data sets using two different starting models. The two independent initial models were created from the initial model obtained in step 3 by randomizing their phases beyond 20Å resolution using EMAN2 (Tang et al., 2007) e2proc3d.py. The data were split in half by dividing each microtubule filament in two halves. For each of the two independent refinements the data was further divided into two halves. These 2 quarter datasets were used to get an estimate of the resolution to which the new reference was filtered for the next refinement cycle. Refinement was done in two stages. The first using Spider (Frank et al., 1996) and the second using Frealign (Grigorieff, 2016). An exhaustive search of the Euler angles and displacements corresponding to each particle-image was performed by multi-reference alignment against projection images of the reference model (Spider operation AP SHC) on 2X binned, CTF corrected particle images (phase flipping only). The orientation and position parameters determined were used to generate a 3D density map by back-projection of the particle-image dataset. Helical symmetry was imposed using himpose (Egelman, 2000). This map was used as the reference model for a new round of multi-reference alignment. Three rounds of the Spider alignment procedure were preformed. The resolution was estimated from the FSC curve calculated using the two 3D maps obtained from the 2 quarters data sets. This resulted in two independently refined 3D maps each with a resolution estimate. After three rounds of alignment the estimated resolution was 6-7 Å. The final cumulative displacement and angular rotations determined in the two Spider refinements were given to Frealign (Grigorieff, 2016) to calculate the two reference models used to perform two independent position/angular refinements and 3D reconstructions. Particle-images were extracted every 82 nm along the filaments (box sizes: 512 × 512 pixels for the NMMT datasets and 616 × 616 pixels for the CTMMT_AMP-PNP_ dataset). To limit the number of overlapping particle-images in each of the two halves datasets used in each of the two independent Frealign refinements the image/particle stack were reordered (and corresponding position angular information) so that odd and even images corresponded to distinct filament quarters. For CTF correction local CTF parameters were estimated with Gctf (Zhang, 2016) (v0.50). Several cycles of Frealign refinement (typically 5) cycles were run limiting the data used to 1/10 Å^−1^, until no further improvement in resolution was detected. Then additional cycles, limiting the refinement data to 1/8 Å^−1^, were run until no further improvement in resolution was detected. At the end of the last cycle, the 2 unfiltered half maps from each independent refinement were averaged to obtain two independently refined maps.

Step 6. Final resolution estimates. Final resolution was estimated using the FSC_0.143_ criteria (Fig. S2) from the two independently refined maps (gold standard) using masks generated with Relion (Scheres, 2012) 2.0 post-process. To make these masks the maps were low-pass filtered to 12 Å, thresholded, the resulting boundaries extended by 2 pixels in all direction and smoothed with a raised cosine edge 5 pixels wide. FSC curves for distinct parts of the maps (microtubule, kinesins or curved tubulin protofilament) were generated by masking the corresponding regions of the map with local masks. These local masks were created by converting to densities the corresponding region atomic models, (EMAN1 (Ludtke et al., 1999) pdb2mrc) low-pass filtering and thresholding them as indicated above. Local resolution was also estimated from the two raw half maps using the Bsoft (Heymann and Belnap, 2007) (v. 1.9.5) function blocres with a kernel size of 10 pixel (Fig. S3).

Step 7. Map post-processing. The two independently refined raw half maps were averaged, corrected for the modulation transfer function of the K2 summit detector and sharpened to restore high resolution contrast with Relion (Scheres, 2012) 2.0 post-processing program. The B-factors used were: −130, −100 and −20 Å^2^ respectively for the NMMT_apo_, CTMMT_AMP-PNP_ and NMMT_AMP-PNP_ maps. The maps were then locally filtered using Bsoft blocfilt according to the local resolution estimates given by Bsoft blocres (step 6). Illustration of the 3D density maps and molecular models were prepared using UCSF-Chimera (Pettersen et al., 2004) and Blender.

**2D class averages of CTNM_AMP-PNP_ complex**. Images of ring-like particles of KLP10A NM construct in complex with curved tubulin protofilaments in the presence of AMP-PNP (Fig. S1D) where aligned and classified using Relion (Scheres, 2012) (v 2.0). CTF was estimated using CTFFIND4 (Rohou and Grigorieff, 2015). First 10016 ring-like structures were picked manually from 464 cryo-EM images by marking their centers and masking them with a circular mask of 540 Å. This set of particles was subjected to three rounds of alignment classification and averaging with 100 class averages in each round. Only particles assigned to classes showing distinct tubulin rings with resolved tubulin dimers were kept for the next cycle. In each cycle the coordinates of the particles in the original images were recalculated based on the position of the average-image particle. This was done by applying the inverse transformation used to align the particles and any additional displacement necessary to re-center the particle in the class average-images. The updated coordinates were used to generate a new set of particles for the next round of alignment classification and averaging. This procedure resulted in three class averages with well centered ring like structures with 15, 14 and 13 tubulin dimers (Fig. 4A). The number of particles in each of these classes were respectively 469, 2171 and 920. We then calculated the coordinates in the original images corresponding to each of two contiguous tubulin dimers in these three class averages. With these coordinate three new sets of particle images (one per each of the three ring-like class average, Fig. 4A) were generated. Each particle-image was masked with a circular mask to included only two tubulin heterodimers (diameter = 176 Å). These three data sets were subjected to three rounds of alignment classification and averaging done as before (centering the particles on the inter-dimer interface of the two tubulin heterodimers approximately at the location where the tip of KLP10A loop-2 would contact tubulin). This procedure resulted in three independently aligned groups of class average images. Two distinct types of class average images could be recognized in each of the three groups. One in which each of the densities corresponding to the two tubulin heterodimers has one kinesin motor-domain density associated and the other with only one kinesin-motor-domain associated density (Fig. 4B). The number of particles associated to each of these two class-average images for each of the three groups (15, 14 and 13 tubulin heterodimers ring like structures) were respectively (954, 1441), (610, 567) and (1109, 530) with the first number in parenthesis corresponding to the class with two motor-domains bound and the second number corresponding to the class with one motor domain bound.

**Model building.** Atomic models of the cryo-EM density maps were built as follow. First, atomic models for each of the proteins chains involved in the complex were generated from their amino-acid sequence by homology modeling using Modeller (Fiser and Šali, 2003). Second, models of the complexes were built by manually placing and fitting the polypeptide chain models into the cryo-EM electron density maps using UCSF-Chimera (Pettersen et al., 2004). At this step N- or C-terminal residues of the chain models falling outside the electron densities (i.e. with no associated density) were deleted from the models. Third, the complex models were flexible fitted into the electron density maps using Rosetta for cryo-EM (DiMaio et al., 2009; Wang et al., 2015). For this step we first generated models using the relax-protocols and picked the models with the best scores (best matching to the cryo-EM density among the lowest 20% energy models). From these models we then generated over 100 models using the iterative-local-rebuilding protocols and picked the ones with the best scores as before. Fourth, we refined the best scored Rosetta models against the cryo-EM density maps using Phenix (Adams et al., 2010) real space refinement tools. Fifth, the Phenix refined models were edited in Coot (Emsley and Cowtan, 2004) and UCSF-Chimera (Pettersen et al., 2004) to resolve a few atomic clashes, geometry problems and areas of the model outside the electron densities. The edited models were then refined again using Phenix (Adams et al., 2010) (step 4). Several iterations of steps 4 and 5 were performed to reach the final atomic models.

## Acknowledgments

We thank The Albert Einstein Analytical Imaging Facility for electron microscopy support; the Albert Einstein College of Medicine High Performance Computing Facility for computing support; Dr. Juan D. Diaz-Valencia for help in the initial parts of the project; Dr. Gary J. Gerfen for critical reading of the manuscript. This work was supported by NIH grant R 01GM113164 (HS).

Cryo-EM data collection was performed at the Simons Electron Microscopy Center and National Resource for Automated Molecular Microscopy located at the New York Structural Biology Center, supported by grants from the Simons Foundation (349247), NYSTAR, and the NIH National Institute of General Medical Sciences (GM103310) with additional support from Agouron Institute [Grant Number: F00316] and NIH S10 OD019994-01. We thank Ed Eng and Bill Rice for assistance during data collection.

Atomic structure models and corresponding cryo-electron densities have been deposited in the protein and electron microscopy data bases (PDB, EMDB) with the following accession codes: PDB ID XXXX, EMD-XXXX; PDB ID XXXX EMD-XXXX; PDB ID XXXX, EMD-XXXX.

## Supplementary Figures

**Fig. S1.**
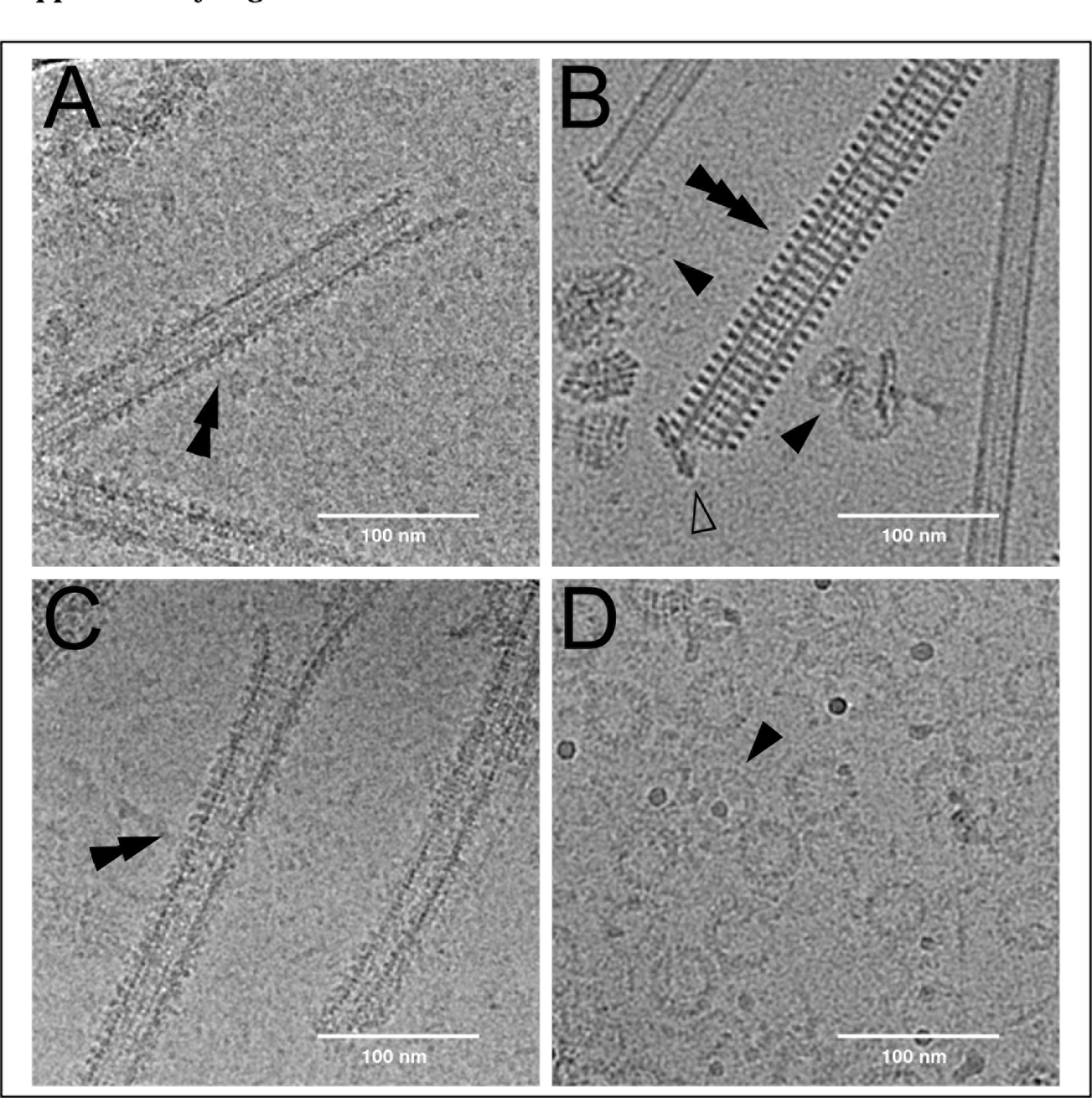
Cryo-electron micrographs. Four examples of cryo-EM images (average of motion corrected movie frames). Each image corresponds to the experimental conditions used to obtain particular datasets: **(A)** NMMT_apo_ complex. **(B)** CTMMT_AMP-PNP_ complex. **(C)** NMMT_AMP-PNP_. **(D)** CTNM_AMP-PNP_. Double arrows in A and C show examples of microtubules with KLP10A decorated lattice. Single arrows in B and D show examples of curved-tubulin-protofilaments-KLP10A complexes. The triple arrow shows a microtubule wrapped around with curved-tubulin-KLP10A complex spirals. The open arrow shows the spiral peeling ways from the microtubule end.

**Fig. S2.**
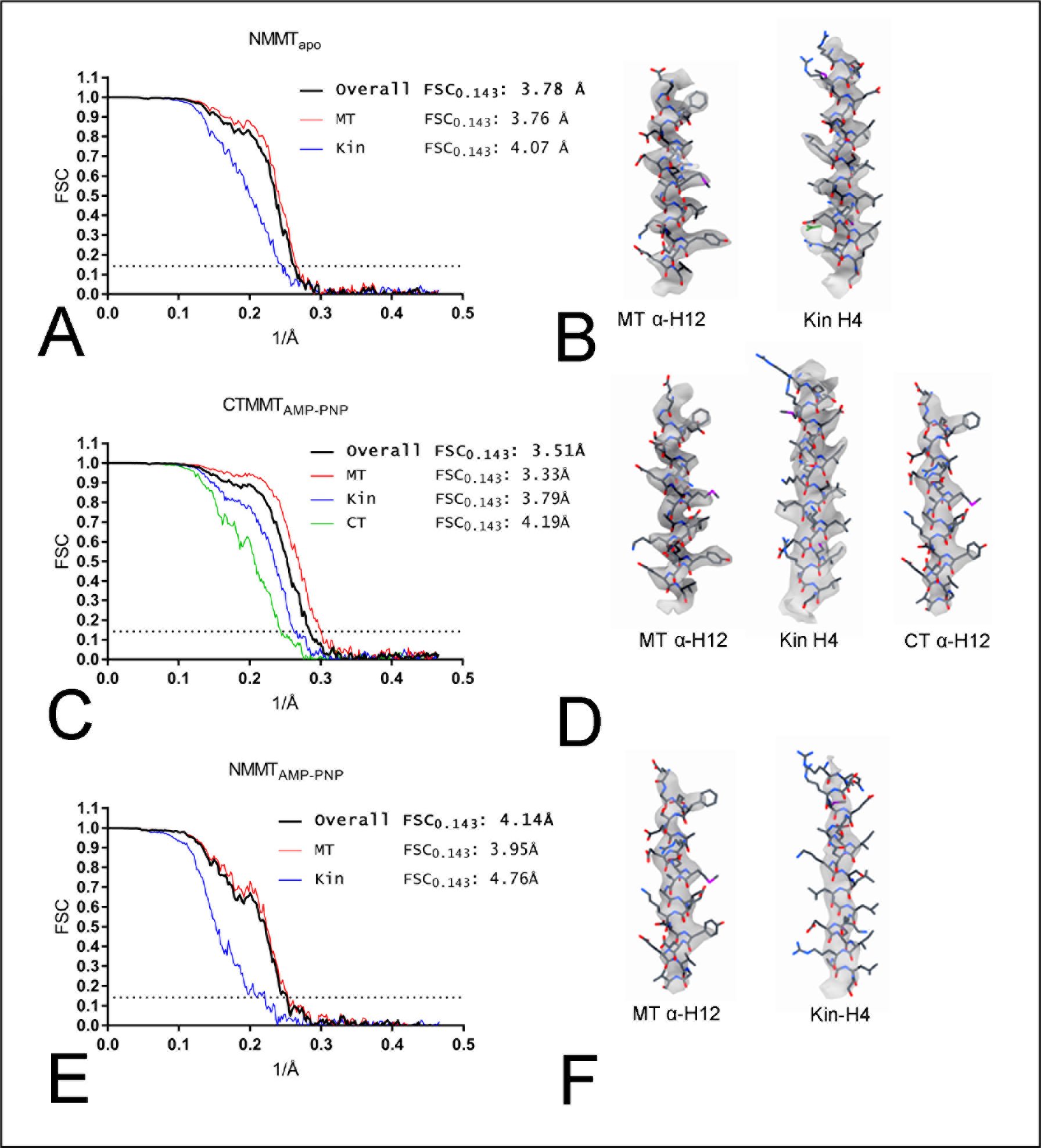
Resolution estimates. Gold standard Fourier shell correlation (FSC) of the 3D reconstructions (left) and examples of the electron densities and corresponding atomic models of some α-helices in the complexes (right). **(A-B)** NMMT_apo_. **(C-D)** CTMMT_AMP-PNP_. **(E-F)** NMMTAMP-PNP. FSC curves corresponding to the whole complex (overall) or distinct parts of the map (MT: microtubule, Kin: KLP10A, CT: curved tubulin) are shown in different colors as indicated. The FSC_0.143_ level is indicated by the dotted line.

**Fig. S3.**
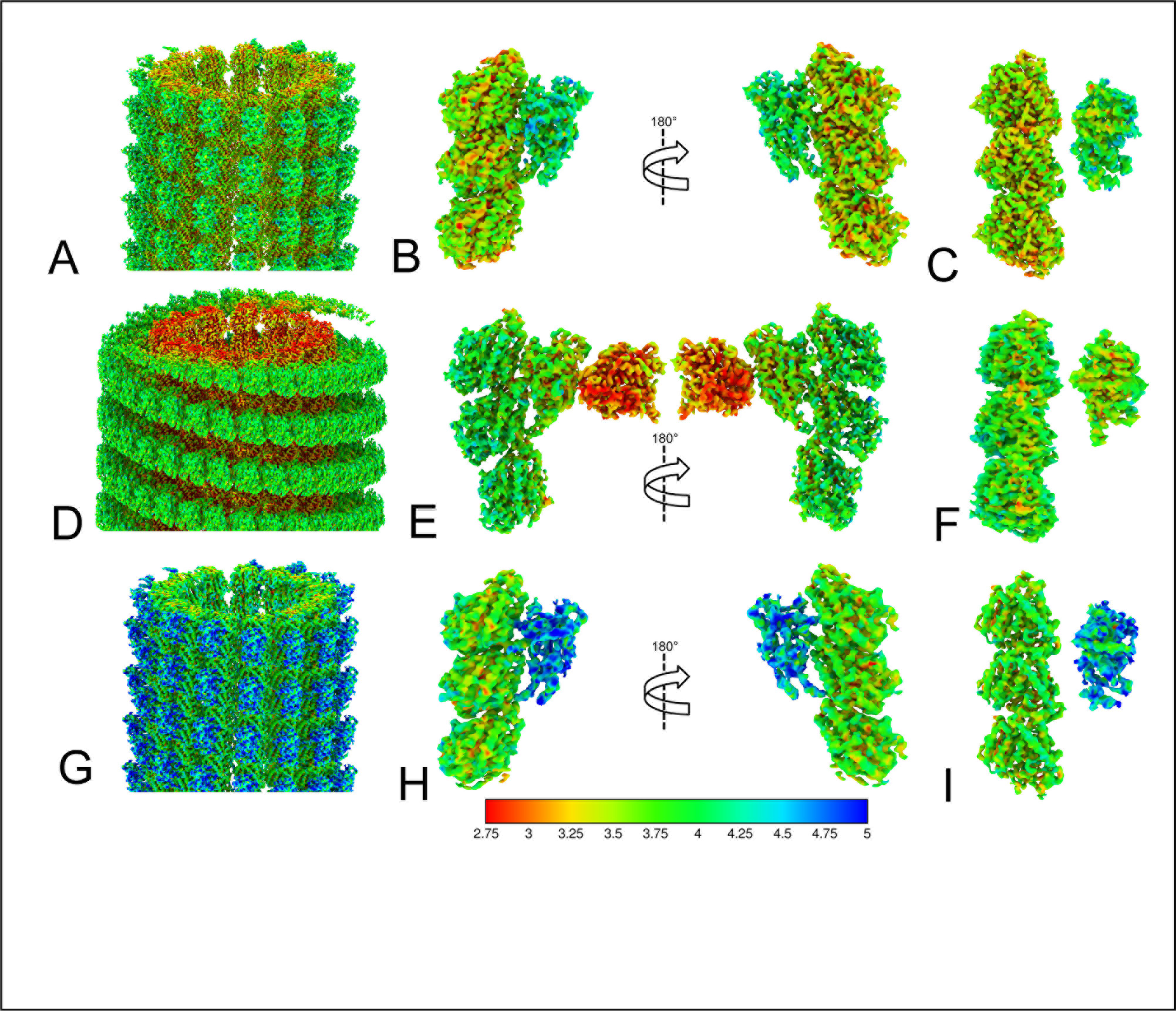
Local resolution maps. .**(A-C)** NMMT_apo_. **(D-F)** CTMMT_AMP-PNP_. **(G-I)** NMMT_AMP-PNP_. Each rows shows: the whole map (left, A, D and G); an asymmetric unit with an extra β-tubulin subunit in two orientations (middle, B, E and H); an asymmetric unit with an extra β-tubulin subunit showing the interface between KLP10A and tubulin (right, C, F, and I). Local resolution was estimated using Bsoft (see Methods). Resolution color scale values in Å.

**Fig. S4.**
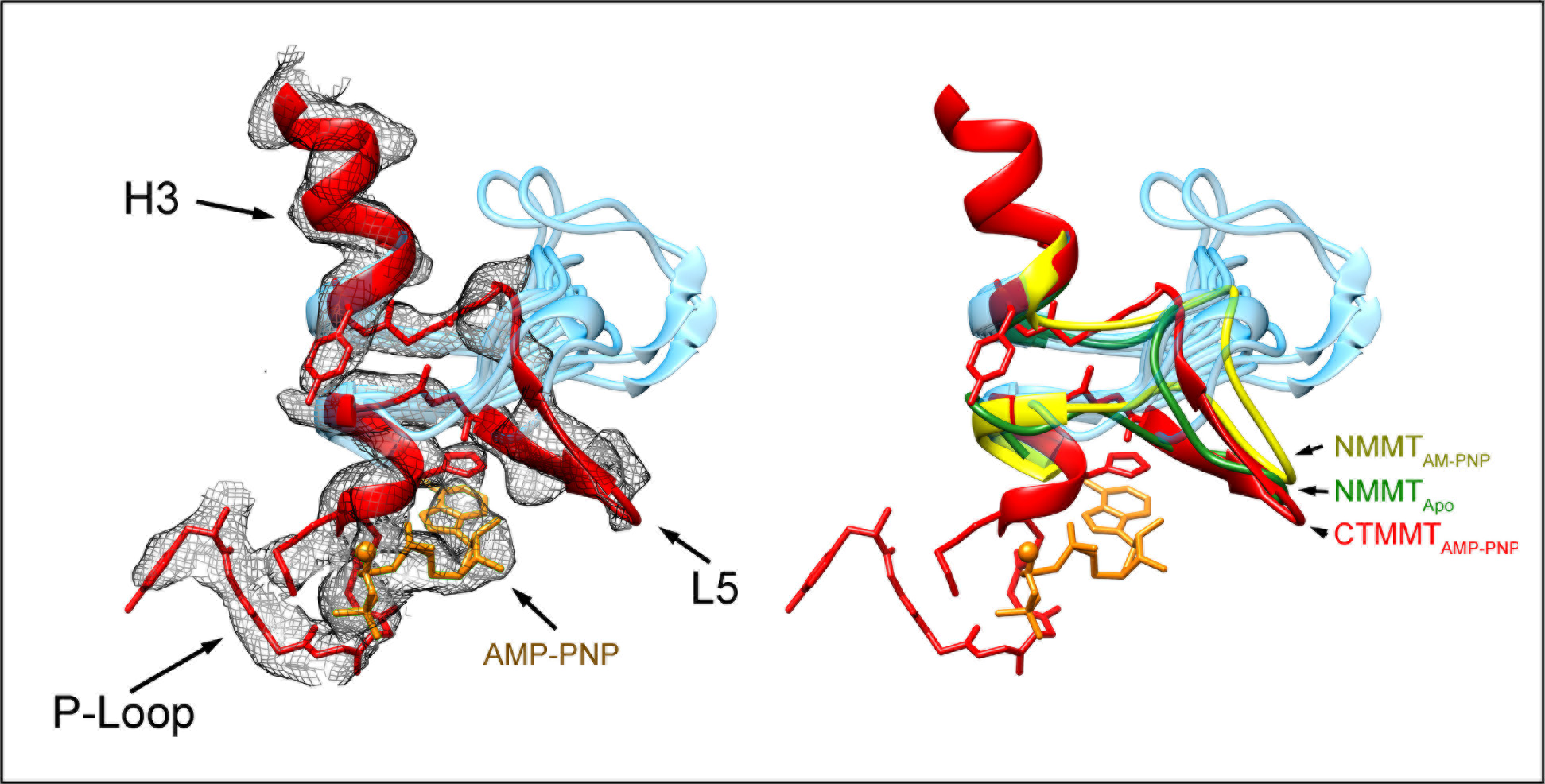
Loop-5. The mesh shows the electron density in the helix-3 (H3) loop-5 (L5) region of the CTMMT_AMP-PNP_ complex with the corresponding atomic model in red and the AMP-PNP molecule in orange. The structure of the L5 of other kinesins motor domains, after aligning the corresponding H3 and P-loop regions, are shown in light blue (PDB accession codes: 1BG2, 4HNA, 4LNU, 1MKJ, 5LT0, 5LT1, 5LT2, 5LT3, 5LT4, 3J8X, 3J8Y, 1VFW, 4OZQ, 3ZFD, 3HQD, 1Q0B, 5X3E, 3LRE, 1RY6, 2NCD). The position and orientation of L5 in the NMMT complexes are relatively similar to the one of the CTMMT_AMP-PNP_ complex (right panel).

**Fig. S5.**
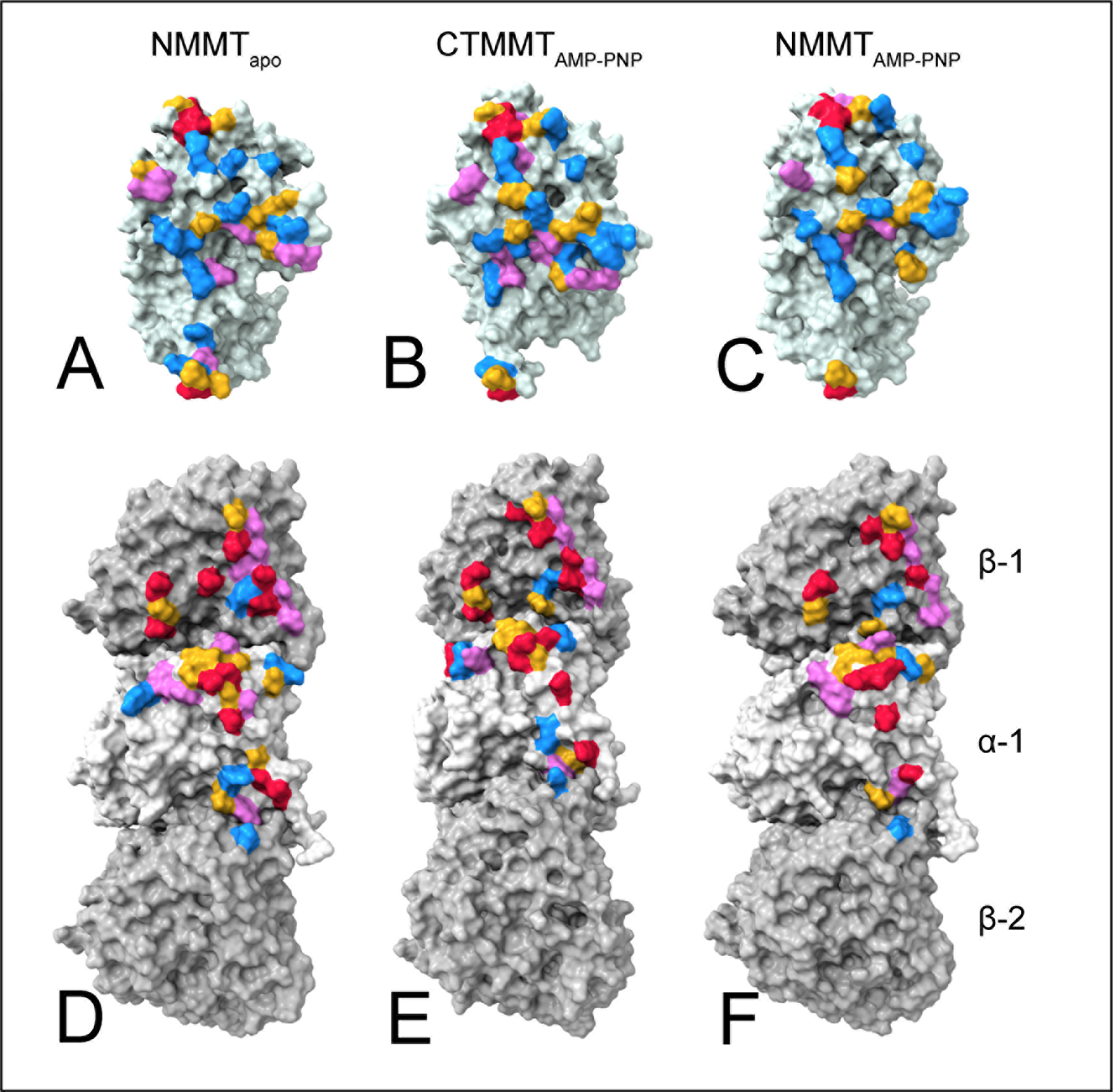
KLP10A-tubulin interface. The figure shows the tubulin and KLP10A complementary interfaces as solvent exclusion surfaces for the three 3D complex models. **(A, D)** NMMT_apo_. **(B, E)** CTMMT_AMP-PNP_. **(C, F)** NMMT_AMP-PNP_. The top row shows the KLP10A side of the interface and the bottom the tubulin side. Residues at the interface that are close enough for contact are colored according to type: negative red, positive blue, polar magenta and non-polar yellow.

**Fig. S6.**
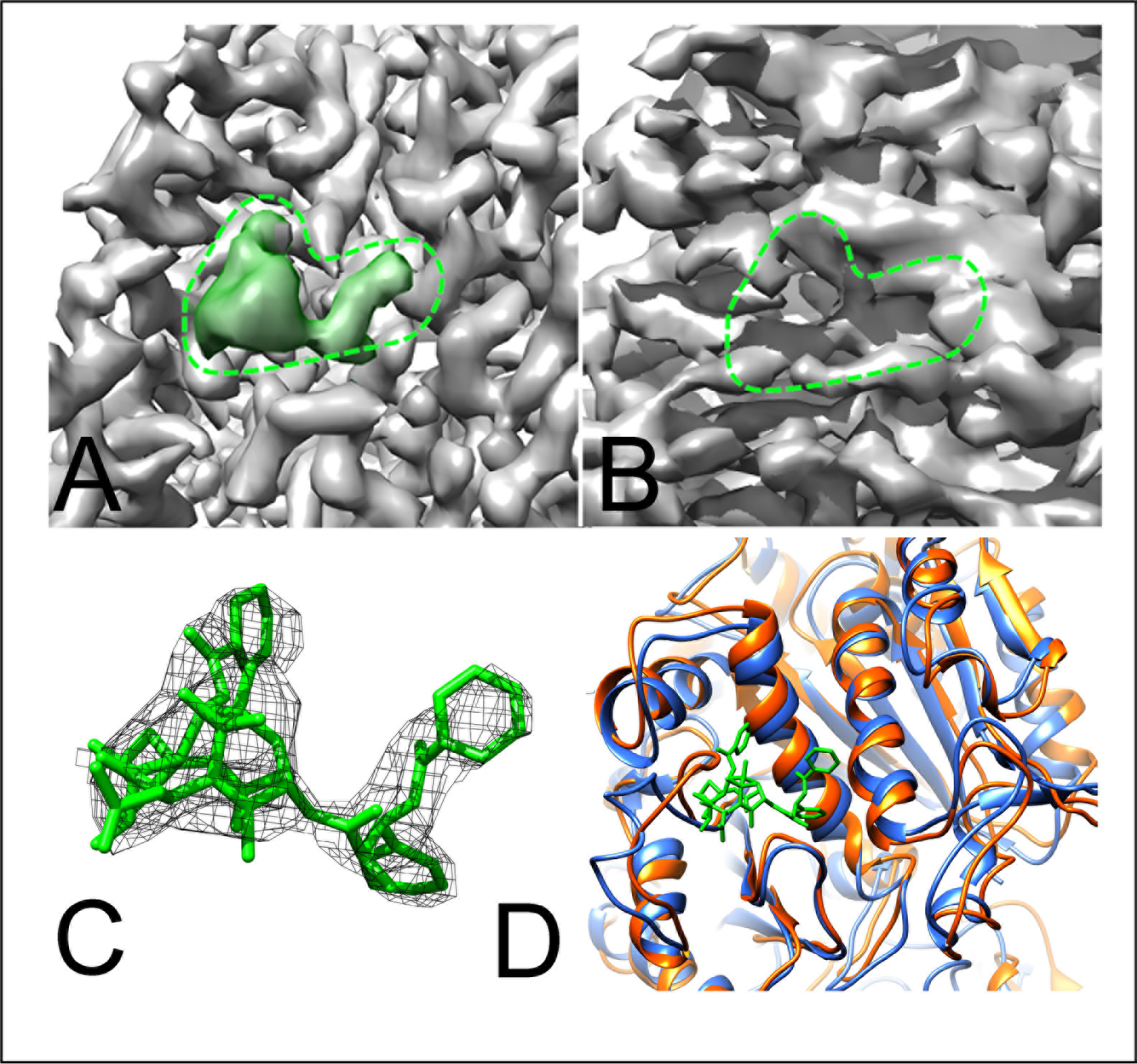
Paclitaxel binding site. **(A)** Cryo-EM density (iso-surface representation) near the paclitaxel (Taxol®) binding site in the β-tubulin subunit of the microtubule in the CTMMT_AMP-PNP_ complex. The density corresponding to the paclitaxel molecule is colored green. The paclitaxel molecule with associate density is showed in C. **(B)** Cryo-EM density near the paclitaxel binding site in the β-tubulin subunit of the curved tubulin (CT) in the CTMMT_AMP-PNP_ complex. Despite the maps having different resolution in this area (Figs. S2 and S3) a distinct cavity can be observed in B in the place that is occupied by paclitaxel in A (dotted green outline). **(C)** Paclitaxel molecule (green) and associated cryo-EM density (dark mesh) in the β-tubulin subunit of the microtubule in the CTMMT_AMP-PNP_ complex. **(D)** Comparison of the two β-tubulin models in the CTMMT_AMP-PNP_ complex. The microtubule β-tubulin is shown in blue with paclitaxel molecule in green. The curved tubulin protofilament β-tubulin is shown in orange.

## Supplemental Movies

**Movie S1.** Morph between NMMT_apo_ and CTMMT_AMP-PNP_ models with structures aligned on the KLP10A P-loop. KLP10A in blue; KLP10A P-loop in pink; KLP10A SW1 in green; KLP10A SW2-H4 in magenta, α-tubulin in light gray, β-tubulin in dark gray.

**Movie S2.** Morph between NMMT_AMP-PNP_ and CTMMT_AMP-PNP_ models with structures aligned on the KLP10A P-loop. KLP10A in blue; KLP10A P-loop in pink; KLP10A SW1 in green; KLP10A SW2-H4 in magenta, α-tubulin in light gray, β-tubulin in dark gray.

**Movie S3.** Morph between NMMTapo and NMMTAMp-pNp models with structures aligned on the KLP10A P-loop. KLP10A in blue; KLP10A P-loop in pink; KLP10A SW1 in green; KLP10A SW2-H4 in magenta, α-tubulin in light gray, β-tubulin in dark gray.

**Movie S4.** Morph between NMMT_apo_ and CTMMT_AMP-PNP_ models with structures aligned on β-tubulin-1. KLP10A in blue; KLP10A P-loop in red; KLP10A SW1 in green; KLP10A SW2-H4 in magenta, α-tubulin in light gray, β-tubulin in dark gray. Nucleotide not shown.

**Movie S5.** Morph between NMMT_AMP-PNP_ and CTMMT_AMP-PNP_ models with structures aligned on β-tubulin-1. KLP10A in blue; KLP10A P-loop in red; KLP10A SW1 in green; KLP10A SW2-H4 in magenta, α-tubulin in light gray, β-tubulin in dark gray. Nucleotide not shown.

**Movie S6.** Morph between NMMT_apo_ and NMMT_AMP-PNP_ models with structures aligned on β-tubulin-1. KLP10A in blue; KLP10A P-loop in red; KLP10A SW1 in green; KLP10A SW2-H4 in magenta, α-tubulin in light gray, β-tubulin in dark gray. Nucleotide not shown.

